# AHR activation accelerates the resolution of TGF-β1 induced fibroblast activation and promotes alveolar type 1 cell regeneration in alveolar organoids

**DOI:** 10.1101/2024.04.29.590972

**Authors:** Andrew S. Hagan, Scott Williams, Casey J. N. Mathison, Shanshan Yan, Bao Nguyen, Glenn C. Federe, Guray Kuzu, Joseph C. Siefert, Janice Hampton, Victor Chichkov, S. Whitney Barnes, Frederick J. King, Brandon Taylor, John R. Walker, Rui Zhao, Jimmy Elliott, Dean P. Phillips, Bin Fang, Rebekah S. Decker

## Abstract

Regeneration of the alveolar epithelium is necessary to restore tissue architecture and gas exchange capabilities in chronic pulmonary diseases such as fibrosing interstitial lung disease. While it is known alveolar type 2 (AT2) cells give rise to alveolar type 1 (AT1) cells to repair the alveolar epithelium after injury, methods to promote this process under pathological settings are poorly understood. Here, using a complex 3D organoid culture with TGF-β1 dependent impaired AT1 spheroid formation, we performed a high-throughput screen (HTS) with ∼16,800 compounds to identify small molecules that increase number of AT1 spheroids. Longitudinal single cell RNA sequencing (scRNA-seq) revealed that DB-11-BE87 increased AT1 regeneration by reducing TGF-β1 induced fibroblast activation, concurrently with AHR activation in those cells. These studies highlight a novel HTS system to identify factors that can promote AT1 differentiation and suggest AHR activation as a method to counteract pathological TGF-β1 signaling in pulmonary disease.

## INTRODUCTION

Efficient air-gas exchange is the fundamental function of the lung and occurs at the alveolar epithelium, the most distal end of the respiratory tree. The alveolar epithelium consists of the squamous alveolar type 1 (AT1) and the cuboidal alveolar type 2 (AT2) cells. Composing fewer absolute numbers than AT2s, AT1s comprise 95% of the alveolar surface due to their elongated morphology.^1^ During homeostasis and injury AT2 cells self-renew and give rise to differentiated AT1 cells to maintain the alveolar epithelium.^2^ Recent work has uncovered transitional cell states along the AT2 to AT1 differentiation trajectory that accumulate during models of pulmonary fibrosis and may have a pathogenic role in disease.^3–5^ Published single cell atlases of idiopathic pulmonary fibrosis (IPF) report a decrease in both AT2 and AT1 cells, confirming a depleted alveolar epithelium in diseased lung tissue.^6,7^ Increasing AT2 to AT1 differentiation and limiting the accumulation of transitional cell states could provide benefit to patients with fibrosing interstitial lung disease.

Alveolar niche fibroblasts populations under homeostasis support AT2 progenitor cell self-renewal.^8^ Conversion of homeostatic fibroblasts to myofibroblasts marks progression of lung fibrosis and is a major contributor of excess extracellular matrix (ECM) deposition in fibrotic lung disease.^9,10^ Intermediate fibroblast to myofibroblast cell states are non-invasive, however they may respond to TGF-β1 stimulation to produce an invasive myofibroblast subpopulation with high ECM expression in the fibrotic lung.^11,12^ TGF-β1 stimulation also inhibits alveolar spheroid formation through fibroblast activation suggesting a decreased ability to support lung epithelial repair.^13^ Decreasing pathogenic fibroblasts subtypes could improve epithelial niche support.

In the mouse, AT1s are marked by *Hopx* expression.^14,15^ For this study, we developed a high-content imaging (HCI) assay to quantify AT1-containing spheroids (*Hopx-GFP*+) derived from a *murine* lung homogenate 3D organoid culture. From a 16,800 compound high throughput screen (HTS), we identified LA-13-ZS70 as a small molecule capable of promoting Hopx-GFP spheroid formation. Single-cell RNA sequencing (scRNA-seq) of DB-11-BE87, a medicinal chemistry refinement of LA-13-ZS70, treated organoids revealed compound dependent promotion of epithelial organoid formation with decreased aberrant transitional AT2-AT1 epithelial cells. Differentially expressed genes (DEGs) and cell-cell interaction analyses suggested aryl hydrocarbon receptor (AHR) in the mesenchyme as a likely target of DB-11-BE87. This was further supported by activation of the AHR pathway in reporter assays. DB-11-BE87 treatment reduced TGF-β1 induced fibroblast to myofibroblast transition, decreasing myofibroblasts and increasing fibroblast transitional states. This work suggests AHR activation in the pulmonary mesenchyme as a beneficial strategy to reduce fibroblast to myofibroblast transition and increase beneficial epithelial niche support.

## RESULTS

### Development of a murine *Hopx-GFP*^+^ assay

To develop a HCI assay for HTS, the alveolar organoid assay initially described by *Barkauskas et al.* was modified.^2^ Briefly, all distal lung cells from *Hopx-GFP* mice were isolated and seeded in optically clear 384-well plates in a mixture of culture medium and 50% Matrigel. After Matrigel solidification, media was added and replaced on day 4 or day 5. At endpoint (∼day 12), spheroid cultures were imaged live to measure *Hopx-*GFP+ spheroids (Fig. 1A). Addition of TGF-β1 impaired spheroid formation in a dose dependent manner, while addition of galunisertib, a TGF-βR1 inhibitor, eliminated this effect and increased spheroid number (Fig. 1B,C). galunisertib promoted spheroid formation in the absence of exogenous TGF-β1 suggesting endogenous TGFβ signaling in culture conditions. To identify molecules that reverted the impaired AT1 spheroid formation, a screen of ∼16,800 compounds was performed on the alveolar spheroid assay in the presence of TGF-β1 (Fig. 1D).^16^ Positive (30µM galunisertib, Fig. 1D, red) and negative (empty well, Fig 1D, blue) controls displayed distinct effects on spheroid formation. 100 hits were called by being >3 standard deviations from median values by both *Hopx-*GFP+ spheroid number and total integrated signal intensity (Fig. 1D, green). LA-13-ZS70 (Fig. 1D, right panel, star) was selected as a candidate for additional validation, and its activity was indicated by concentration-response curve (EC_50_ = 376 nM, count of *Hopx-GFP*+ spheroids) (Fig. 2A,B). To confirm an increase of markers of AT1 cells we performed a highly multiplexed gene expression readout with RNA-mediated oligonucleotide annealing, selection, and ligation with next-generation sequencing (RASL-seq).^17^ We assembled a panel of alveolar genes by literature mining for canonical cell lineage markers and sorted AT1 cells (*Hopx*-GFP+, Sftpc-tdTomato-, CD31-, CD45-) vs. AT2 cells (*Hopx*-GFP-, *Sftpc*-tdTomato+, CD31-, CD45-) for AT1 and AT2 specific markers (Supp. Fig. 1; Table 1). Higher expression of AT1 lineage markers, including *Ager*, *Aqp5*, and *Gprc5a*, and airway signaling pathway genes were observed upon treatment with LA-13-ZS70 compared to the DMSO control, confirming the improved alveolar organoid regeneration in the presence of LA-13-ZS70 (Fig. 2G, Table 2).

**Figure 1:**
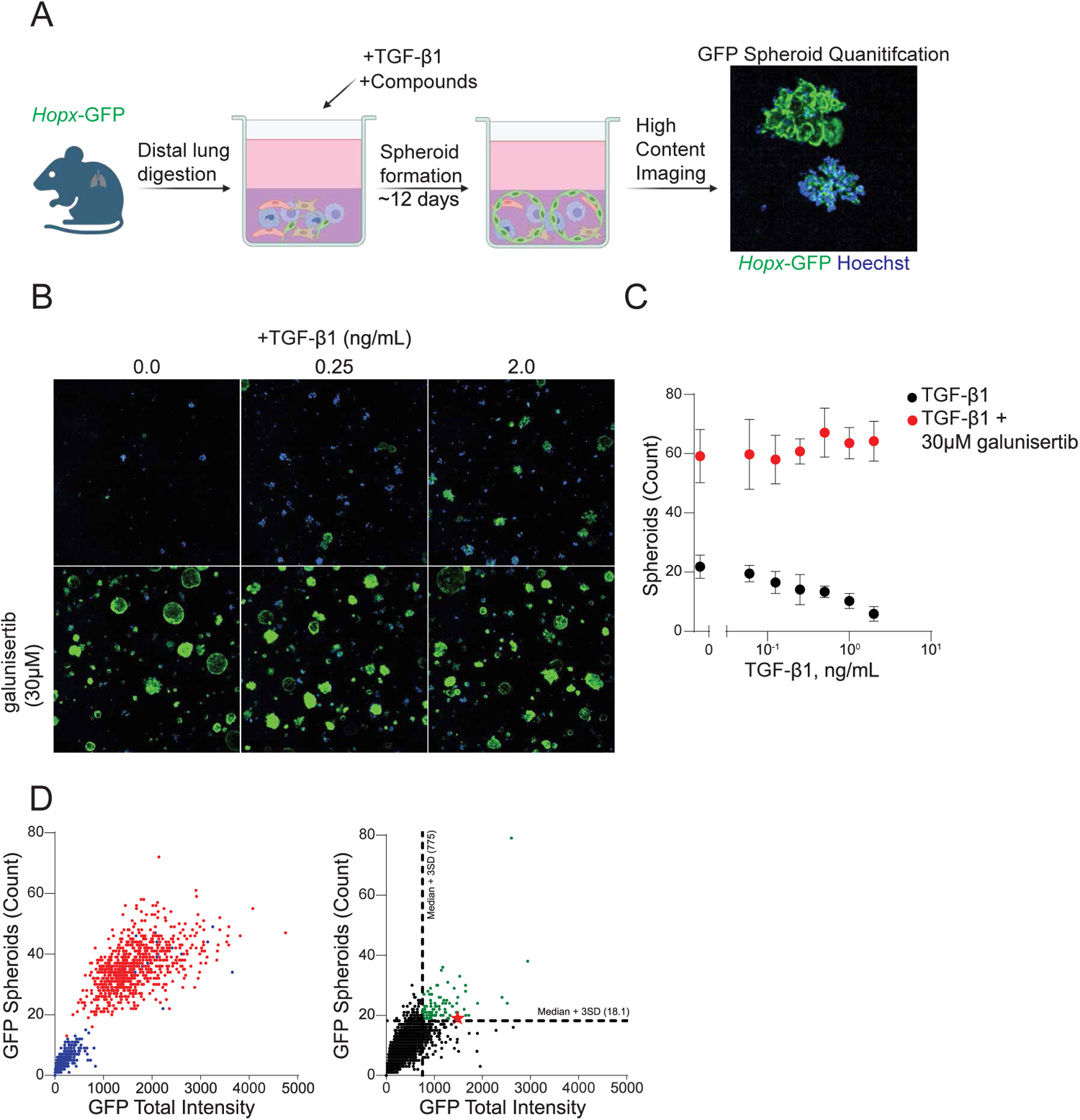
High-throughput screening on multi-cellular murine alveolosphere assay identifies small molecules that rescue TGFβ impaired alveolar type 1 spheroid formation. **A**: Schematic of assay. Whole distal lung cells from *Hopx-GFP* mice were plated in 384-well plates, cultured in Matrigel for ∼12 days to promote spheroid formation, and imaged live for GFP signal to quantify alveolar type 1 spheroids. **B**: Representative images of live alveolospheres treated at time of plating with increasing doses of TGF-β1 with or without 30 µM of TGF-βR1 inhibitor galunisertib. **C**: Quantification of B. n=12, mean ± SD. **D**: Quantification of GFP spheroids vs. GFP total intensity for controls (left) or small molecule library (right). Red = positive control (30 µM galunisertib). Blue = negative control (no compound addition). Green = called compounds with biological activity. Black= compound without called biological activity. Star = LA-13-ZS70; primary hit for followup (right panel). Dotted line indicates Median +3SD.

**Figure 2:**
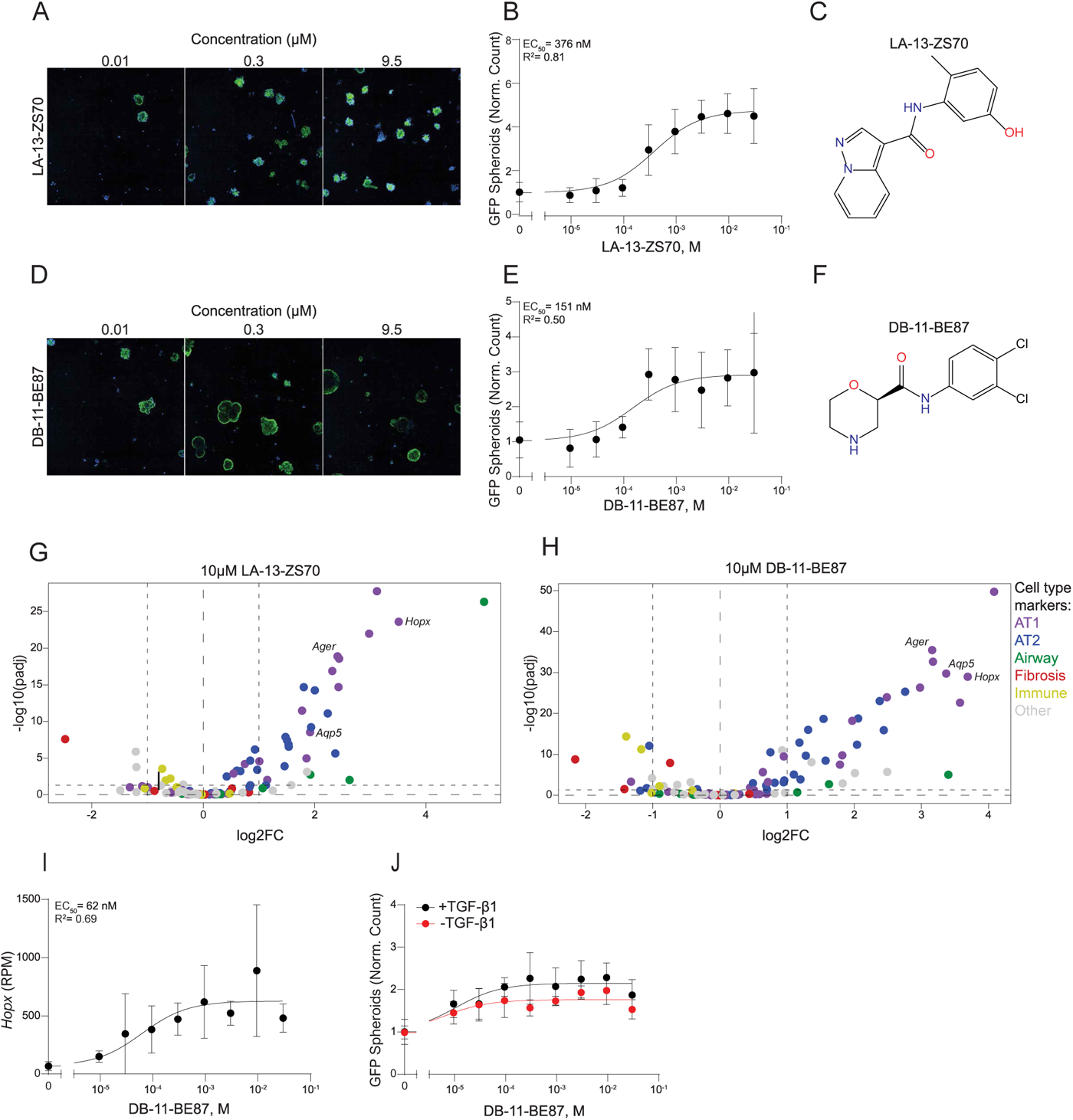
DB-11-BE87 increases AT1-AT2 differentiation with partial dependency on TGFB pathway activation. **A**: Representative images of live alveolospheres treated at time of plating with increasing doses of LA-13-ZS70. **B**: Quantification of concentration-response curve for LA-13-ZS70 on GFP adjusted count spheroids. EC_50_= 376 nM, R^2^=0.81. n=4 for all concentration except untreated well, n=96 for untreated well. mean ± SD **C**: Chemical structure of LA-13-ZS70. **D**: Representative images of live alveolospheres treated at time of plating with increasing doses of DB-11-BE87. **E**: Quantification of concentration-response curve for DB-11-BE87 on GFP adjusted count spheroids. EC_50_= 151 nM, R^2^=0.50. n=4 for all concentration except untreated well, n=96 for untreated well. mean ± SD **F**: Chemical structure of DB-11-BE87. **G**: RASL-seq of live alveolospheres treated at time of plating with 10 µM LA-13-ZS70. Dotted line indicated ± gene fold change of 2 and p-adjusted value of 0.05 compared to DMSO control. Plotted as mean ± SD. Purple =AT1 genes; Blue= AT2 genes; Green= Airway genes; Yellow=fibrosis genes; Red= Immune genes; Gray=other genes. n=4 **H**: RASL-seq of live alveolospheres treated at time of plating with 10 µM DB-11-BE87. Dotted line indicated ± gene fold change of 2 and p-adjusted value of 0.05 compared to DMSO control. Plotted as mean ± SD. Purple =AT1 genes; Blue= AT2 genes; Green= Airway genes; Yellow=fibrosis genes; Red= Immune genes; Gray= other genes. n=4 **I**: Quantification of concentration-response curve for DB-11-BE87 on *Hopx* expression. EC_50_= 62 nM, R^2^=0.69. n=3-5 for all concentration except untreated well, n=129 for untreated well. mean ± SD **J**: Quantification of concentration-response curve for DB-11-BE87 on GFP adjusted count spheroids in culture with or without exogenous TGF-β1. + TGF-β1 EC_50_= 10.5 pM, R^2^=0.70. - TGF-β1 EC_50_= 6.3 pM, R^2^=0.65. n=6 for all concentration except untreated well, n=24 for untreated wells. mean ± SD

Although LA-13-ZS70 improved spheroid formation, it displayed poly-pharmacology with a multi-kinase inhibition profile (BTK, EPHA4, EPHB4, KIT, LCK, MAPK14, PDGFRA; data not shown). Subsequent medicinal chemistry efforts informed by structure-activity relationship generated DB-11-BE87, a structural analogue of LA-13-ZS70 (Fig. 2C,F). A concentration-response curve confirmed activity of DB-11-BE87 (DB-11-BE87: EC50 = 151 nM, count of *Hopx-GFP*+ spheroids) (Fig. 2D,E). In addition, significant increases (log2 fold change >1 and FDR <0.05) of canonical AT1 markers including *Ager*, *Aqp5, Gprc5a*, *Hopx*, and *Igfbp2* were observed by RASL-seq (Fig. 2H; Table 2). Hopx expression displayed a similar concentration-response curve to the Hopx-GFP+ imaging readout in DB-11-BE87 treated samples (Fig 2.E,I). Activity of DB-11-BE87 was attenuated when TGF-β1 was removed from culture suggesting dependency (Fig. 2J). Together, our HTS identified DB-11-BE87 that reverted TGF-β1 impaired AT1 spheroid formation.

### Longitudinal scRNA-seq of alveolar organoid culture identifies pulmonary fibrosis associated *Fn1*+ aberrant epithelial cells

To understand the effect of DB-11-BE87 on mixed distal lung cell cultures, longitudinal scRNA-seq was performed at three different time points (day 1, 3, and 11) following TGF-β1 and compound treatment (DMSO, 10 µM DB-11-BE87, 10 µM QA-92-TQ17) (n=3-4 per group) (Fig. 3A). Compound QA-92-TQ17 was chosen as a minimally active structural analogue of the active DB-11-BE87 (Supp. Fig. 2). As a control, cells freshly dissociated from untreated mouse lung on day 0 were profiled by scRNA-seq. After QC, 117,450 cells were obtained from all conditions. Cells from four major categories were identified using canonical markers and were clearly separated on uniform manifold approximation projection (UMAP), a dimensional-reduction technique for improved visualization and interpretation of single cell data (Fig. 3A).^18^

**Figure 3:**
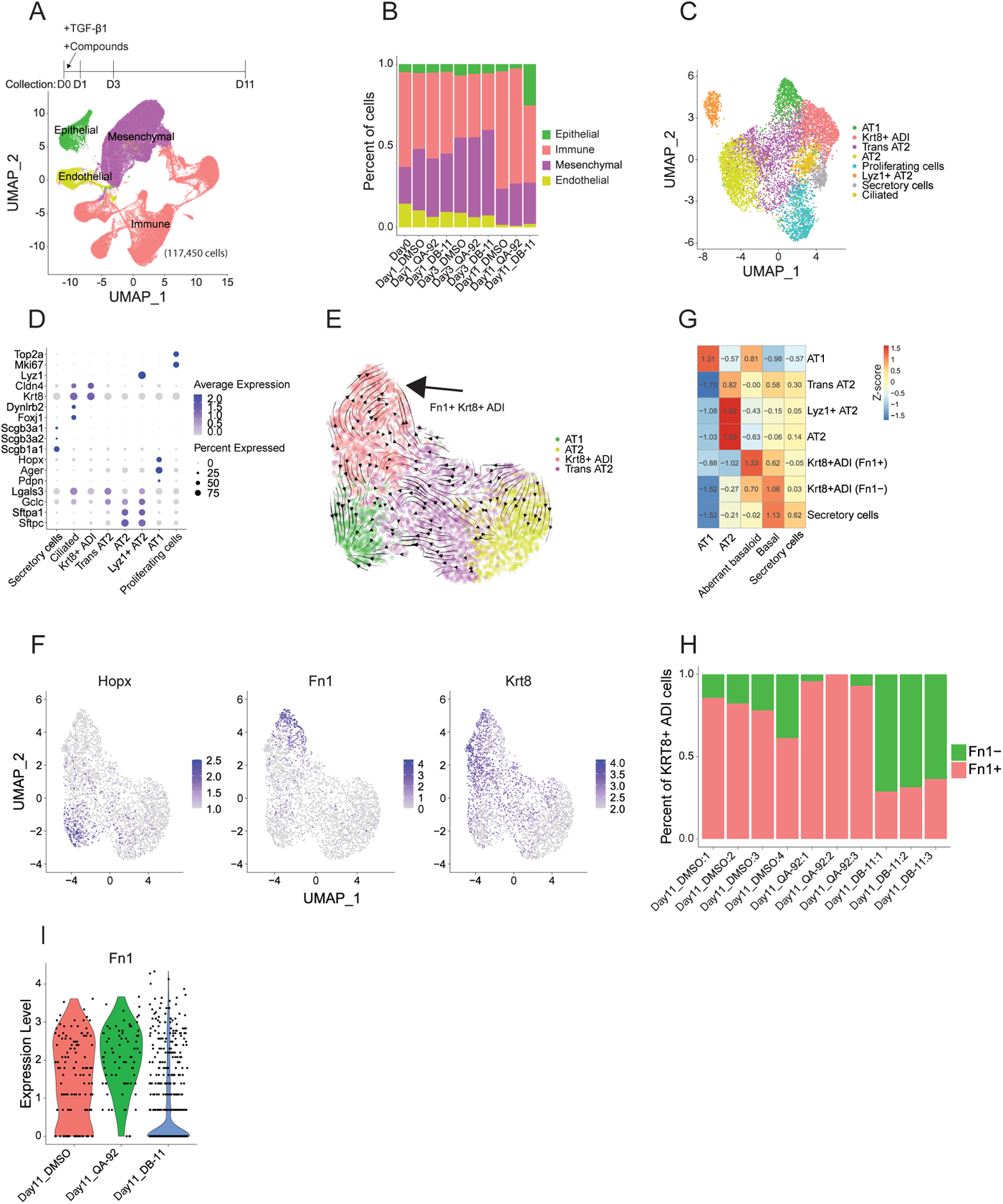
Alveolosphere scRNA-seq detects *Fn1*+ aberrant epithelial cells. **A**: Single cell suspension from alveolosphere assay were analyzed using scRNA-seq at indicated timepoints post assay setup. Untreated control (Day 0) and three treatment groups (DMSO, 10 µM DB-11-BE87, 10 µM QA-92-TQ17) were included to understand DB-11-BE87’s influence on TGF-β1 treated alveolosphere culture. 4 cell categories containing 117,450 integrated cells from all samples are colored on UMAP. **B**: Cell proportions of major cell categories for the indicated timepoint and compound treatment in the alveolosphere assay. **C**: UMAP for all epithelial cells integrated from all conditions. 8 cell types and states were identified. **D**: Expression of canonical epithelial cell type markers. **E**: RNA velocity vectors projected over UMAP plot of AT1, AT2, *Krt8*+ ADI, and Transitional AT2 epithelial cell types, from all samples. **F**: Expression of individual genes among cells from E. **G**: Degree of similarity between indicated cell types/states from patient derived samples (x-axis) and alveolosphere culture (y-axis). Color code indicates scaled spearman correlation coefficient (z-score) of two transcriptomes. **H**: Fraction of *Krt8*+ ADI cells that expressed *Fn1,* across individual samples collected on day 11. **I**: Violin plot of *Fn1* expression in *Krt8*+ ADI cells from samples in three treatment groups.

The proportion of the four cell categories in the cell culture changed over time, although significant differences in the composition of compound-treated samples and the DMSO controls were not observed at day 1 or day 3. By day 11, there was a marked increase in the population of epithelial cells in the DB-11-BE87-treated sample (p=2.6e-6, student’s t-test Fig. 3B); consistent with the increased *Hopx-GFP* observed in the HCI assay. To examine epithelial cell types in our samples, we integrated epithelial cells from each condition and identified cell clusters based on the transcriptome similarity. Cell type labels were assigned to each cluster according to previously established cell type markers (Secretory cells: *Scgb3a1*, *Scgb3a2*, *Scgb1a1;* Ciliated: *Dynlrb2*, *Foxj1*; AT2: Sftpa1, *Sftpc*; Lyz1+ AT2: *Lyz1*; AT1: *Ager*, *Hopx*, *Pdpn*; Proliferating cells: *Top2a*, *Mki67*).^3,4^ Notably, clusters resembling recently described transitional AT2-AT1 cell states that persist in mouse fibrosis models were captured in this *in vitro* assay (Trans *AT2: Lgals3* high, *Gclc* high; *Krt8*+ alveolar differentiation intermediate (ADI) : *Krt8* high, *Cldn4* high) (Fig. 2C,D).^3,4^

To determine lineage relationship between these populations, the ratio of spliced to unspliced transcriptomic reads was calculated to deduce the RNA velocities and predict differentiation trajectories across cells.^19^ As expected, RNA velocity trajectory indicated a clear differentiation of AT2 cells into trans AT2 and then AT1 cells (Fig. 3E; Supp. Fig. 3A). Interestingly, the trajectory also suggests a bifurcate fate of *Krt8*+ ADI cells, one part differentiated into AT1 while the rest transitioned into another terminal cell state featured by high expression of *Fn1* (hereafter *Fn1*^+^ *Krt8*^+^ ADI, Fig. 3E,F). A similar result was also observed by diffusion pseudo-time analysis (Supp. Fig. 3B,C).^20^ Recent scRNA-seq studies identified aberrant basaloid cells, also known as KRT5^-^/KRT17^+^ epithelial cells, in the distal lung of IPF patients, which exhibited high level of *Fn1*.^6,7^ To examine if the *Fn1*^+^ *Krt8*^+^ ADI cells in TGF-β1 treated mouse lung cell organoids are similar to the aberrant basaloid cell population in IPF patients, the transcriptomes of alveolar epithelial cells from the two species were compared. High similarity was observed between the *Fn1*^+^ *Krt8*^+^ ADI and the aberrant basaloid cells (Fig. 3G). Interestingly, the DB-11-BE87 treatment significantly reduced the population of *Fn1^+^* cells among *Krt8*^+^ ADI cells (Fig. 3H, student’s t-test p=0.002), and resulted in lowered *Fn1* expression in the culture on day 11 (Fig. 3I). Together, our data suggest that TGF-β1 treated alveolar organoids contain a population similar to pulmonary fibrosis associated *Fn1*+ aberrant epithelial cells, which can be ameliorated by treatment with DB-11-BE87.

### TGF-β1 induces fibroblast activation in alveolar organoids

TGF-β1 is a master regulator that stimulates fibroblast to myofibroblast differentiation and contributes to tissue remodeling in fibrotic disease.^21^ To examine the effect of TGF-β1 on alveolar organoid formation, we inspected temporal transcriptional changes in the mesenchymal compartment of the alveolar organoid culture (Fig. 3A). Alveolar fibroblasts, peribronchial fibroblasts, pericytes, smooth muscle cells, and adventitial fibroblasts were identified in the freshly disassociated cells on day 0 (Supp. Fig. 4A).^11^ One day after growing in media, reduced expression of mesenchymal cell markers, e.g., *Col1a1*, *Col3a1*, *Acta2*, and *Postn*, was observed in all cell clusters (Supp. Fig. 4C). Concurrently, cells became more dispersed on UMAP projection suggesting a rise in transcriptome diversity because of adaption to culture environment (Supp. Fig. 4B).

On day 3, markers of mesenchymal cell types once again became prominent (Fig. 4A,B). Comparing to the cells on day 1, several newly emerging cell populations on day 3 were identified on the integrated UMAP, including clusters 0,5,6,7,8,10 (Fig. 4A,C). Cluster 5,6,7, and 8 showed high level of *Cthrc1* with cluster 5 and 8 also exhibiting a proliferative profile (Fig. 4A,F). By examining the RNA velocity trajectories, cluster 6 and 7 were likely derived from transitioning alveolar fibroblasts and adventitial fibroblasts, respectively (Fig. 4D,E). A recently published atlas of collagen-producing cells in mouse lung has identified a subpopulation of fibroblasts that express *Cthrc1*. This subpopulation emerges in the bleomycin-induced lung fibrosis model and has been found in patients with IPF and scleroderma.^11^ Comparison between the two datasets through integration suggests that *Cthrc1*^+^ fibroblasts observed in the two models are closely similar to each other (Supp. Fig. 5A). Accordant with the reported fibrotic nature of these cells, we observed gene expression profiles consistent with fibrosis-related pathways such as the TGFβ, WNT, and ECM pathways (Fig. 4F).

**Figure 4:**
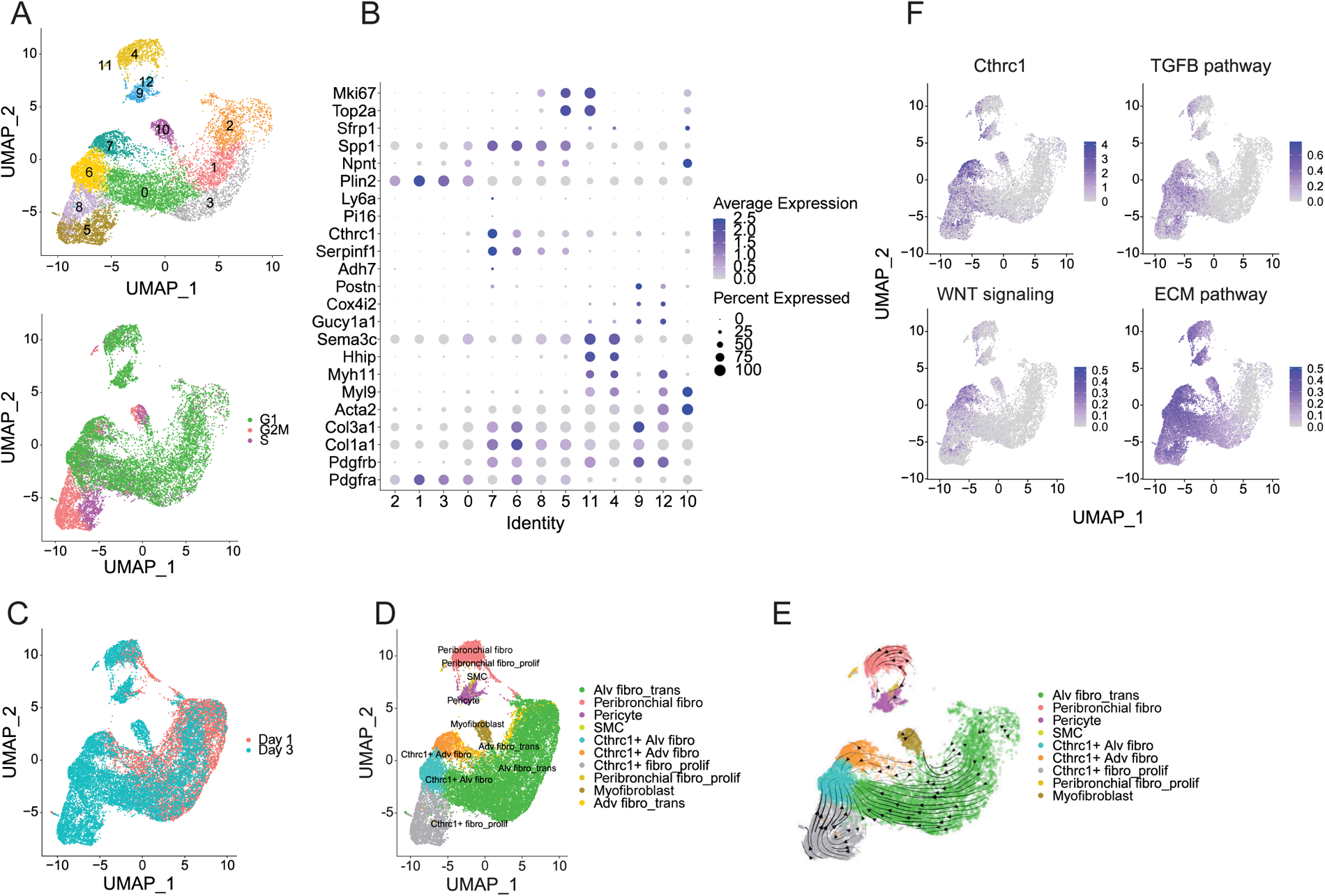
TGFβ induces fibroblast activation in alveolosphere model. **A**: UMAP plots of mesenchymal cell clusters on day 3 (top) and cell cycle phases (bottom, G1: green, G2M: pink, M: purple). Plot contains all 3 treatment groups (DMSO, 10 µM DB-11-BE87, 10 µM QA-92-TQ17). **B**: Expression of marker genes in mesenchymal cell clusters from A. **C**: UMAP plot of integrated mesenchymal cells from day 1 (red) and day 3 (teal). **D**: UMAP plot of mesenchymal subclusters from C colored by cell type labels. **E**: RNA velocity vectors projected over UMAP plot of mesenchymal cells from day 3 samples. Cells are colored by cell type labels. **F**: UMAP plots of mesenchymal cells from day 3 samples, colored by either gene expression (*Cthrc1*) or pathway activity scores (TGF-β pathway, WNT signaling, and ECM pathway).

The newly emerged cell cluster 10 exhibited the highest level of *Acta2* expression and its transcriptomic profile was distinct from smooth muscle cells (Fig. 4B). RNA velocity analysis indicated that they originated from the transitioning alveolar fibroblasts, suggesting a fibroblast to myofibroblast transition already underway by day 3 (Fig. 4E). Interestingly, neither DB-11-BE87 nor QA-92-TQ17 exhibited any influence on this transition, as composition of mesenchymal cells in the organoids was comparable across all treatments (Supp. Fig. 5B).

### DB-11-BE87 resolves TGF-β1 induced fibroblast activation in the alveolosphere model

Mesenchymal cells respond to physical and chemical stimuli to change cellular state.^21^ The integrated UMAP shows that cells from day 11 formed distinct non-overlapping clusters with cells from day 3, suggesting that mesenchymal cells underwent a dramatic shift of cell state by day 11 (Fig. 5A). Most cells from day 11 accumulated into two unique clusters; cluster 3 and 5 (Fig. 5B). Cluster 3 represented myofibroblast cells expressing the highest level of *Acta2* and exhibiting enrichment of muscle contraction and inflammation pathways (Fig. 5C,D,E). Cluster 5 displayed low expression of *Acta2* but high levels of *Sfrp1* and *Spp1*, two recently emerged markers of intermediate states between normal fibroblasts and the *Cthrc1*^+^ pathogenic fibroblasts that accumulate during fibrosis.^12^ In addition, these cells contained highly active adipogenesis genes, exemplified by *Lpl*, implying a lipo-fibroblast identity (Fig. 5C,D,E).

**Figure 5:**
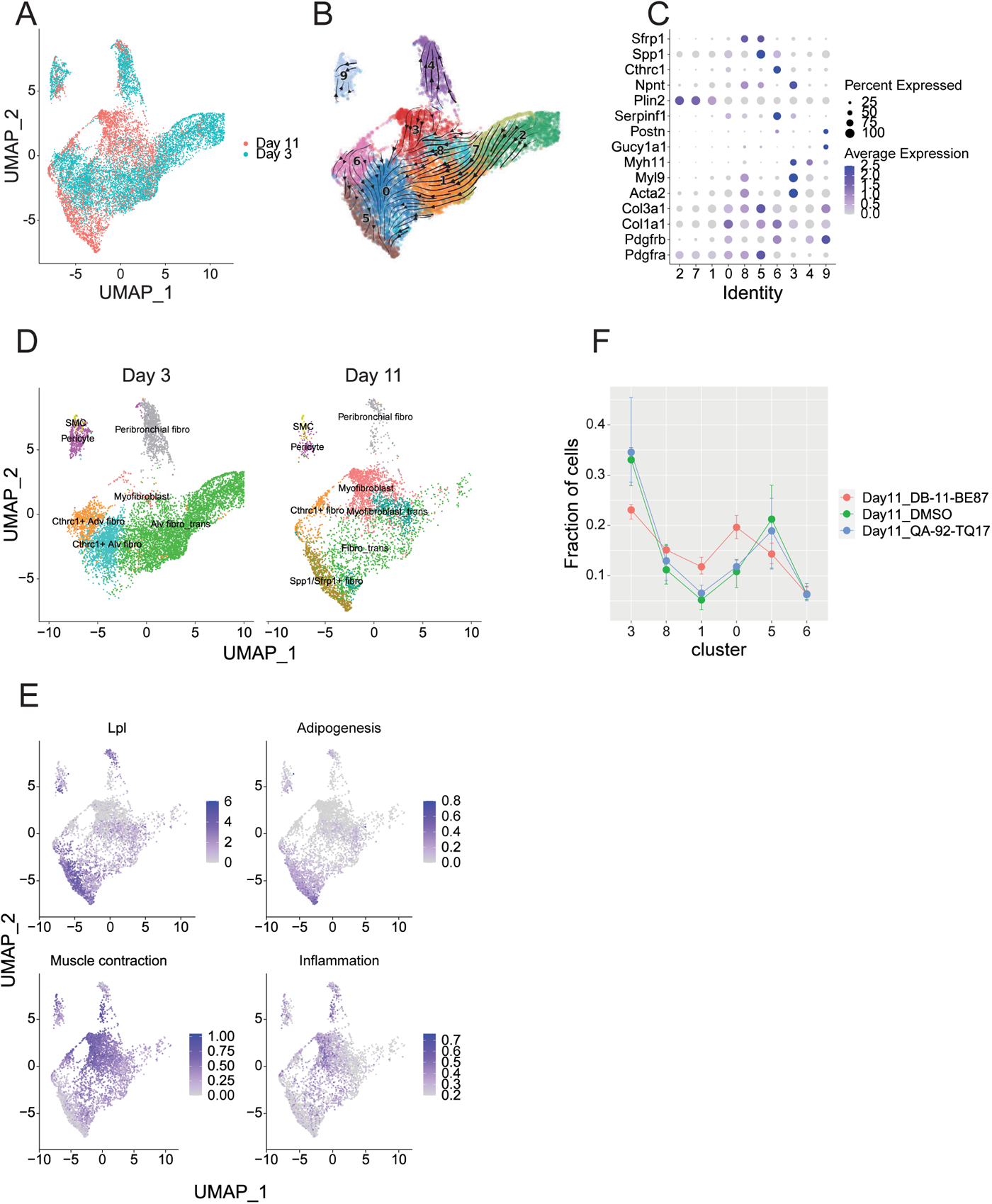
DB-11-BE87 activates AHR and resolves TGF-β1 induced fibrosis in the alveolosphere model. **A**: UMAP plot of integrated mesenchymal cells from day 3 (teal) and day 11 (red). Plot contains all 3 treatment groups (DMSO, 10 µM DB-11-BE87, 10 µM QA-92-TQ17). **B**: RNA velocity vectors projected over UMAP of integrated mesenchymal cells from day 3 and day 11 samples. Cell clusters are numbered. **C**: Expression of marker genes in mesenchymal cell clusters from B. **D**: UMAP plots of mesenchymal cells on day 3 (left) and day 11 (right), colored by cell type labels. **E**: UMAP plots of mesenchymal cells colored by either gene expression (*Lpl*) or pathway activity scores (adipogenesis, muscle contraction, and inflammation), from day 11 samples. **F**: Proportions of mesenchymal cell clusters from B, in day 11 samples separated by compound treatment. * indicates p<0.05 in Student’s t-test; error bar indicates mean ± SD.

To examine the dynamics of the cell type transition, RNA velocity trajectories were projected onto the UMAP (Fig. 5B). Myofibroblast cells appeared to undergo a de-differentiation process, generating transitional cell populations that eventually became the *Spp1^+^/Sfrp1^+^* fibroblasts (Fig. 5B,D). This de-differentiation process implies that TGF-β1 supplementation no longer sustains fibroblast activation on day 11. Notably, a significant acceleration of myofibroblast de-differentiation was observed in the DB-11-BE87 treated samples, leading to a decrease in the proportion of myofibroblast cells and increase in the proportion of transitional fibroblast cells compared to other treatments (Fig. 5F). Together, these data suggest that DB-11-BE87 accelerates the resolution of TGF-β1 induced fibroblast activation in the mouse alveolosphere culture model.

### DB-11-BE87 activates AHR and rescues TGF-β1 impaired AT1 spheroid formation while AHR antagonists do not

To identify the target and mechanism of action (MoA) of DB-11-BE87, we compared DB-11-BE87-treated samples with the DMSO controls and identified differentially expressed genes (DEGs) across unsupervised cell clusters from all four cell categories. To minimize the impact of secondary effects, we investigated cells from the earliest time point (day 1) and identified signaling pathways and biological processes that were enriched in the DEGs from each cell cluster. As expected, most cell clusters did not exhibit significantly perturbed pathways due to the lack of DEGs at early time points. Nevertheless, several mesenchymal cell clusters exhibited shared and significantly perturbed pathways such as Aryl hydrocarbon receptor (AHR) pathway (Supp Fig. 6). By examining genes involved in those pathways, *Cyp1a1* and *Cyp1b1* were the most frequently observed genes up-regulated by DB-11-BE87 (Supp. Fig. 6B). Since *Cyp1a1* and *Cyp1b1* are well-established downstream targets of AHR, we sought to examine the hypothesis that DB-11-BE87 functions as an AHR agonist. A list of canonical AHR downstream targets were assembled, including *Cyp1a1*, *Cyp1b1*, *Cyp2a1*, *Ahrr*, *Tiparp*, *Ptgs1*, *Ptgs2*, and *Nqo1*^22,23^, and module activity of these genes was calculated for each mesenchymal cell cluster (methods). Significant linear correlations were observed between *Ahr* expression and AHR module activity in the DB-11-BE87-treated cells at all time points (Fig. 6A). In contrast, AHR module activities in DMSO controls remained below the background level across cell clusters and time points. Interestingly, although the minimally active analogue QA-92-TQ17 showed weak AHR activation on day 1 and day 3, its effect dropped to the background level on day 11. This is consistent with its minimal effect on AT1 regeneration in the *Hopx-*GFP alveolar organoid imaging assay (Supp. Fig. 2).

**Figure 6:**
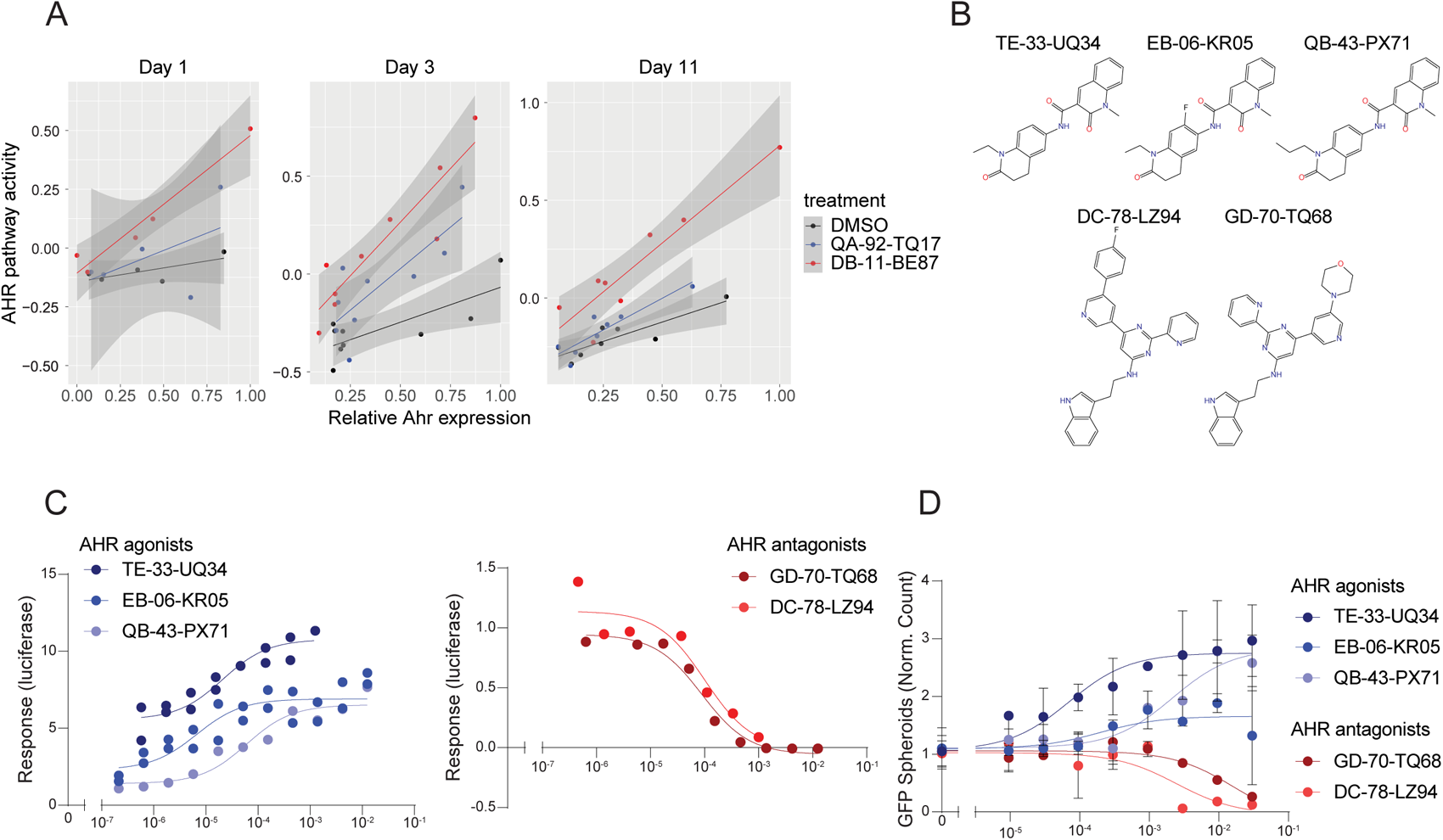
AHR agonists rescue TGF-β1 impaired alveolar type 1 spheroid formation while AHR antagonists do not. **A**: Correlation between Ahr expression and AHR pathway activity across mesenchymal cell clusters from day 1, day 3, and day 11 samples separated by compound treatment. Y-axis indicates average expression of AHR pathway genes relative to random background genes in log scale. **B**: Chemical structure of AHR agonists and antagonists. **C**: Quantification of concentration-response curve for AHR agonists and antagonists on normalized luciferase luminescence in AHR responsive reporter assay. TE-33-UQ34 EC_50_= 21 nM, R^2^=0.90. EB-06-KR05 EC_50_= 6.7 nM, R^2^=0.79. QB-43-PX71 EC_50_= 60 nM, R^2^=0.93. GD-70-TQ68 EC_50_= 89 nM, R^2^=0.98. DC-78-LZ94 EC_50_= 101 nM, R^2^=0.90. Each point is individually plotted. **D**: Quantification of concentration-response curve for AHR agonists and antagonists on GFP adjusted count spheroids in murine alveolosphere *Hopx*-GFP+ spheroid assay. TE-33-UQ34 EC_50_= 70.2 nM, R^2^=0.77. EB-06-KR05 EC_50_= 196 nM, R^2^=0.21. QB-43-PX71 EC_50_= 2.1 μM, R^2^=0.59. GD-70-TQ68 EC_50_= 16.2 μM, R^2^=0.29. DC-78-LZ94 EC_50_= 2.5 μM, R^2^=0.51. n=2 for all concentration except untreated well, n=48 for untreated wells. Some untreated wells are replicated between cmpd curves. mean ± SD.

To confirm effect of AHR activation in the murine *Hopx-*GFP spheroid assay we investigated three AHR agonists (TE-33-UQ34, EB-06-KR05, QB-43-PX71) and two antagonists (GD-70-TQ68, DC-78-LZ94) (Fig. 6B). Compounds were validated as AHR agonist or antagonists based on their demonstrated activity in a murine H1L1.1c2 cell line stably expressing an AHR response luciferase reporter (Fig. 6C). In the murine alveolosphere assay AHR agonists increased GFP adjusted count spheroids while AHR antagonists decreased this count (Fig. 6D). Together these data suggest that DB-11-BE87 resolves TGF-β1 induced fibroblast activation through behaving similar to an AHR agonist.

### DB-11-BE87 induced AT2 to AT1 differentiation through IL-1β signaling

It has been reported that HIF1A-mediated glycolysis controls AT2-AT1 conversion in the bleomycin-injured mouse lung, and that this process is promoted by IL-1β signaling via interstitial macrophages.^5^ In line with this model, we observed that DB-11-BE87 treatment resulted in a shift of cell state in the interstitial macrophages on day 11, towards an activated status featured by high expression of *Il1b* (Fig. 7A,B). Meanwhile, highly active hypoxia and glycolysis pathways were observed in the epithelial cells of the DB-11-BE87-treated organoids (Fig. 7C).

**Figure 7:**
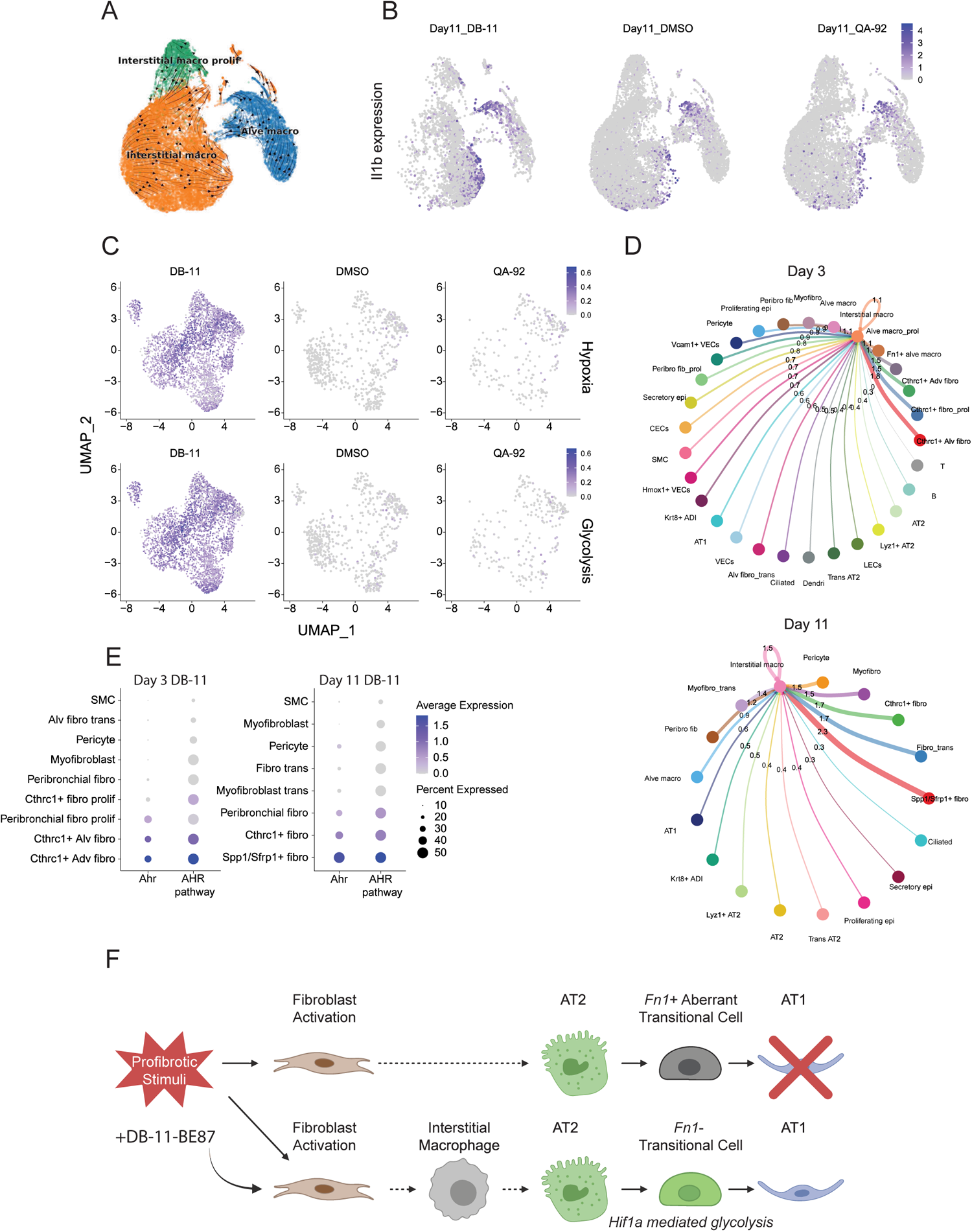
DB-11-BE87 induces hypoxia mediated glycolysis in AT2 on day 11 via IL-1β signaling. **A**: RNA velocity vectors projected over UMAP of macrophage subclusters from day 11 samples. **B**: UMAP plots of macrophage cells colored by *Il1b* expression (purple) from day 11 samples split by compound treatment. **C**: UMAP plots of epithelial cells from day 11 samples, colored by pathway expression scores (hypoxia, top; glycolysis, bottom). Samples split by compound treatment. **D**: Circle plots of cell-cell communications going into interstitial macrophages on day 3 (top) and day 11 (bottom). Thickness of line is proportional to overall communication probability. **E**: Relative expression of *Ahr* and Ahr pathway genes in mesenchymal cell clusters from DB-11-BE87-treated samples on day 3 (left) and day 11 (right). **F**: Schematic presentation of mechanism of DB-11-BE87.

Since DB-11-BE87 activates AHR signaling in mesenchymal cells, we hypothesized that *Il1b* induction in interstitial macrophages was trigged by cell-cell communication between fibroblasts and macrophages. To characterize the niche environment in the organoids, we quantified communication probabilities between cell type pairs using CellChat.^24^ Results show that mesenchymal cells produced the most outgoing signals among four categories, suggesting that they are the primary drivers of the niche environment in the organoids (Supp. Fig. 7A). Interestingly, *Cthrc1+* fibroblasts and *Spp1/Sfrp1*^+^ fibroblasts are the two cell types that generate the most communication signals directed towards interstitial macrophages on day 3 and day 11, respectively (Fig. 7D). This coincides with the strongest *Ahr* expression and AHR module activity in the two cell types in DB-11-BE87-treated organoids (Fig. 7E). These results imply that *Il1b* production in interstitial macrophages is associated with AHR activation in fibroblast cells.

To identify AHR downstream targets that mediate communications between fibroblasts and macrophages, we inspected predicted ligand-receptor pairs between the two *Ahr*-expressing fibroblast subtypes and the interstitial macrophages from day 11 (Supp. Fig. 7B). There are 31 ligands produced in both *Cthrc1+* fibroblasts and *Spp1/Sfrp1*^+^ fibroblasts, mediating potential communications with interstitial macrophages. However, majority of them were also expressed in a wide range of cell types, arguing against their connection to the DB-11-BE87-induced AHR activation (Supp. Fig. 7D). In addition, only one ligand *Sema6d* was commonly up-regulated by DB-11-BE87 in both fibroblast subtypes (Supp. Fig. 7C). These results imply that AHR activation in fibroblasts may not be the direct trigger of *Il1b* induction in the interstitial macrophages. Rather, a more complex communication network among multiple cell types present in the culture may be involved in this process. Together, our study suggests that DB-11-BE87 induces AHR activation in fibrotic fibroblasts, which accelerates the resolution of TGF-β1 induced fibroblast activation and promotes AT2 to AT1 differentiation through an indirect induction of macrophage-mediated IL-1β signaling (Fig. 7F).

## DISCUSSION

Regeneration of the alveolus is a major goal in modern pulmonary biology. Central to alveolar regeneration is re-establishing the correct structure of the damaged epithelium including promoting the differentiation capacity of AT2 to AT1 cells that form the site of air-gas exchange. TGF-β1 and other signaling molecules from pathological mesenchyme direct AT2s into an abnormal intermediate cell state which accumulates in fibrosis.^7,25^ Using a complex multicellular 3D organoid culture with TGF-β1 dependent impaired AT1 spheroid formation, we identified the small molecule DB-11-BE87 as able to rescue spheroid formation. Subsequent scRNA-seq and reporter assays suggests AHR activation in the mesenchyme reverts the deleterious effects of TGF-β1 signaling on AT2-AT1 cell regeneration. This line of inquiry emphasizes the strength of using multicellular cultures coupled with scRNA-seq to identify cell non-autonomous MoAs for early drug discovery.

AHR is a ligand activated transcription factor whose primary function is controlling xenobiotic metabolism.^23^ Following ligand binding, AHR translocates to the nucleus and induces transcription of target genes, including cytochrome p450 enzymes (CYP1A1, CYP1A2, CYP1B1), leading to hydrocarbon detoxication. Generation of *Ahr* null mice identified multiple physiologic roles for AHR outside of exogenous metabolism including regulation of the immune response, hepatic development, regeneration, and cancer.^26,27^ Recent epithelial-specific removal of AHR unveiled its function as an activator of differentiation factors and a repressor of stemness factors during epithelial injury in organoid culture.^28^ The role of AHR in lung epithelial cell regeneration has not been well studied.

AHR controls complex transcriptional events in a ligand, cell-type, and context-specific manner through acting as a sensor for a wide variety of compounds.^29,30^ AHR has been extensively studied in the context of activation by polycyclic aromatic hydrocarbons found in cigarette smoke and immunomodulation in respiratory disease.^31,32^ Additional evidence suggest AHR modulates tissue fibrosis ^33^. In the murine model of bleomycin induced pulmonary fibrosis, removal of AHR has a heightened fibrotic response while activation of AHR has an attenuated response.^34,35^ Our results suggest a benefit to AHR activation in these rodent models could be through promoting AT1 cell regeneration, a characteristic of recovery after bleomycin injury.^36^ While currently approved drugs for idiopathic pulmonary fibrosis (Nintedanib and Pirfernidone) target fibroblast activation, it is unclear if these therapies promote AT1 cell regeneration suggesting AHR activation could have novelty.

Acquisition of a contractile and extracellular matrix synthesizing fibroblast phenotype is a defining feature of fibrosing interstitial lung disease.^37,38^ TGF-β1 is the dominant perturbation known to trigger this phenotypic conversion and has been studied extensively in fibrosis for over the past 30 years.^39^ TGF-β1 treated human fibroblasts have impaired ability to support mouse AT2 spheroid formation.^13^ AHR ligands decrease expression of *Tgfb1* and inhibit TGF-β1 induced myofibroblast formation.^40–42^ Crosstalk between the TGFβ and AHR pathways has been described, although the physiological importance of this interaction is not completely understood.^43,44^ Our results suggest that AHR activation in fibroblasts reverts the pathogenic effects of TGF-β1 and restores the innate epithelial support role of the alveolar stroma.

Target identification and MoA studies following on hits from phenotypic screening is notoriously difficult.^45^ Target identification is desired to define on-target limitations, design confirmatory follow-up experiments, and identify pharmacodynamic markers for efficacy *in vivo*.^46^ Lack of target identification has historically led to discontinuation of drug discovery programs hampering progression of promising molecules to the clinic. Here we leveraged scRNA-seq to identify the mesenchyme as the primary site of action of DB-11-BE87, identifying a cell-non autonomous effect of AHR activation on promoting alveolar regeneration. These insights would have been un-realized with traditional bulk transcriptomic approaches and highlight the need for understanding phenotypic screens at a single-cell resolution.

This study leveraged multiple technical approaches to identify a new MoA in AT1 regeneration. HTS with a HCI readout in a multicellular assay was used to probe a diverse chemical library to find modulators of a desired phenotype. RASLseq was performed to confirm hits and to inform subsequent medicinal chemistry. scRNA-seq was executed for MoA determination. We propose these approaches are replicable and present an avenue for identifying new MoAs in forward pharmacology.

### Limitations

There are interspecies differences between mouse and humans’ ability to respond to AHR ligands.^47^ We have no evidence to suggest AHR activity is beneficial for human alveolar regeneration. DB-11-BE87 increases AHR target gene expression however it is unclear if DB-11-BE87 directly binds to AHR acting as a ligand or if it activates the AHR pathway though a disparate mechanism. While DB-11-BE87 increases the AHR pathway activation, the dependence of AHR for DB-11-BE87’s efficacy has not been established. Further work on testing the effect of DB-11-BE87 on human alveolar cultures, direct binding to AHR, and efficacy dependence of AHR will extend the clinical relevance of this work.

## MATERIALS and METHODS

### Mice

All mice (*Mus musculus*) were housed in a barrier facility free of pathogens. All studies were performed under a protocol approved by the Institutional Animal Care and Use Committee at Novartis-San Diego (Protocol No. 20-429). Mouse strain *Hopx^3FlagGFP^*, referred to as *Hopx-GFP*, has been described.^48^ All mouse strain (*Hopx^3FlagGFP^*, referred to as *Hopx-GFP*; Sftpc*^CreER^*; *ROSA26^tdTomato^*) have been described.^36,48,49^

### *Hopx-GFP* murine organoid assay

Adult *Hopx-*GFP+ mice were sacrificed with isoflurane, perfused with PBS through the right ventricle, and the lungs were extracted. Trachea, connective tissue, and upper bronchiole was removed under dissection microscope. Lungs were cut into small pieces (>2 mm^2^) and incubated in protease solution for 30 minutes at 37°C in a shaking incubator. Protease solution contained: 450 U/mL Collagenase Type I (Gibco; 17100– 017), 4 U/mL Elastase (Worthington Biochemical Company; LS002279), Dispase 5 U/mL (BD Biosciences; 354235), and 0.33 U/mL DNaseI (Roche; 10104159001) dissolved in DMEM/F12 (Gibco; 11320033). Digestion enzymes were inactivated with DMEM/F12 + 10% FBS, cells were centrifuged at 500*g* and resuspended in Trypsin/DNAse solution and incubated for 30 minutes at 37°C in a shaking incubator. Trypsin/DNase solution contained: 0.1% Trypsin-EDTA (Gibco, 25200056) + 0.13 U/mL DNaseI (Roche; 10104159001) in DMEM/F12. Trypsin/DNase solution was inactivated with DMEM/F12 + 10% FBS, cells were filtered through a 70 µM cell strainer, centrifuged at 300g (2x), resuspended in RBC cell lysis buffer (Sigma; R7757), and digestion enzymes inactivated with DMEM/F12 + 10% FBS. Cells were centrifuged at 500*g*, resuspended in DMEM/F12 + 2% FBS, counted on a hemacytometer, and diluted to 1 x 10∧^6^ cells per mL. Cells were filtered through a 40 µM cell strainer, resuspended 1:1 in Matrigel Growth Factor Reduced (Corning; 354230), and 20 µL of solution (10,000 cells) was plated into the bottom of 384-well CellCarrier Ultra tissue culture plate (PerkinElmer; 6057300) pre-spotted with compounds (10 µM final concentration). 30 µM of TGFβ-R1 inhibitor galunisertib was pre-spotted as a positive control. Once solidified 50 µL of ITS media was added per well and plates were incubated in a standard cell culture incubator. ITS media contained: DMEM/F12 + 5% FBS, 1/100 ITS-G (Gibco; 41400045), 1/100 Antibiotic-Antimycotic (Gibco; 15240096), 0.5 ng/mL TGFβ1 (R&D Systems; 240-B-002). Fresh ITS media was added on top after 4-5 days in culture. On day 8 60µL of ITS media was removed and replaced with 60µL of fresh ITS media. On day 10-11 ITS media was removed and fresh ITS media containing a 1:2500 dilution of Lysoview-405 (Biotium; 70066) was added. Plate was placed in cell culture incubator for ∼24 hrs before live fluorescent imaging for *Hopx^GFP^* was performed on high content confocal imaging platform: ImageXpress Micro (Molecular Devices; primary screen) or Opera Phenix (PerkinElmer; reconfirmation and subsequent experiments).

### HCI Imaging, Processing, and Data Normalization

The image z-stacks of the *Hopx-*GFP channel were compressed to maximum intensity projections (MIPs). Representative *Hopx-*GFP MIPs were then used to train an ilastik pixel classifier to differentiate between spheroids, artifacts, dim objects, and background.^50^ The spheroid probability images output from ilastik together with MIPs from *Hopx-*GFP channel were passed to a CellProfiler pipeline for object segmentation and numeric feature measurements.^51^ Spheroids were initially identified by thresholding the spheroid probability images (> 0.5). Spheroid morphological and intensity features were then measured, and small (< 500 pixels) and dim (*Hopx-*GFP upper quartile intensity < 0.0075) spheroids were filtered out. Mean aggregation of the spheroid features was utilized to generate well-level data, was written to CSV, and analyzed in Spotfire (Tibco). For adjusted measurements data was normalized by setting negative control wells to 1.

### Small molecule compound deck

Primary small molecule screening deck consisted of ∼16,800 compounds selected based on diversity and attractive chemical properties (molecular weight, cLogP, solubility, etc).^16^

### Single cell sample prep

*Hopx*-GFP murine organoid assay was established similar to above except scaled to plate 4-7 drops of 50 µL cells-medium/Matrigel mixture in each well of a 6-well plate. At designated times (day 0, day 1, day 3, day 11) cells were removed from Matrigel for scRNA-seq. Briefly, media was removed and washed with ice cold PBS. TrypLE Express (Gibco; 12604013) was added to wells and incubated at 37°C, followed by scraping of Matrigel bubbles to collect in a 15 mL conical tube. Cells were mechanically disrupted by pipetting ∼10 times and incubated at 37°C for an additional 10 minutes. Cells were mechanically disrupted again and enzyme was inactivated with DMEM/F12 + 10% FBS + 1/100 Penicillin-Streptomycin (Gibco; 15140122). Cells were centrifuged at 300*g* at 4°C, resuspended to 1000 cells/ µL in DMEM/F12 + 10% FBS, and filtered through a Flowmi 40 µM cell strainer (Sigma; BAH136800040).

### AHR agonist and antagonist assay

H1L1 1c2 cells stably expressing a dioxin response element (DRE) driven luciferase reporter were plated at 5,000 cells/well in 40uL of medium. Medium contained MEMa + 10% FBS, 1% Antibiotic-Antimycotic, 1% Non-essential amino acids, 1% HEPES, 1% Sodium pyruvate, and 200 μg/mL geneticin. Compounds were added at indicated molarity. Plates were centrifuged for 15 seconds at 1000rpm and mixed with a plate mixer. Plates were placed in cell culture incubator for 18 hrs. 20μL of luciferase reporter gene assay reagents was added (Bright Glo, Promega, E2610) and luminescence was on a multimodal plate reader. For antagonist assay 3 nM of dioxin was added at time of cell plating.

### Single cell transcriptomics

The Chromium Next GEM Single Cell 3’ reagent kit v3.1 from 10X Genomics (protocol CG000204 Rev C) was used in scRNA-seq library preparation. Single-cells were partitioned into nanoliter-scale Gel Beads-in-emulsion (GEMs) using Chromium Controller. Upon cell lysis, the poly-A tails of mRNA are captured on the beads containing an Illumina read 1 sequencing primer, 16nt 10X barcode, 12nt UMI and 30nt poly (dT) sequence. Reverse transcription occurs followed by amplification of full-length cDNA from polyA mRNA. Enzymatic fragmentation and size-selection was used to optimize cDNA amplicon size. End repair, A-tailing, Adaptor ligation and PCR create the final sequencing libraries that contain the P5 and P7 primers used in Illumina bridge amplification. The libraries were sequenced using NextSeq 550 High Output Kit v2.5 (150 cycles).

### Flow sorting AT2 and AT1 populations

Adult Sftpc*^CreER^; ROSA26^tdTomato^; Hopx-*GFP mice were sacrificed with pentobarbital, perfused with PBS through the right ventricle, and the lungs were instilled with 1.5mL of enzymatic cocktail. Enzymatic cocktail contained: 5 U/mL Dispase (Worthington; NPRO2), 4 U/mL Elastase (Worthington; 2294), 200 U/mL Collagenase Type 1 (Gibco, 10104159001), 0.33 U/mL DNaseI (Roche; 10104159001) in PBS. Lung was removed from chest and incubated in an additional 3.5mL of enzymatic cocktail for 45 minutes at room temperature. Lung was transferred to dish containing 10mL of DNase I solution and pulled apart completely using forceps. DNase I solution contained: 1 mg DNase I in 10mL DMEM (11965118) + 10% FBS + 1% Penicillin-Streptomycin (Gibco; 15140122). Lung tissue in DNase I solution was gently rocked at low speed for 10 minutes at room temperature. Cell suspension was filtered through a 100 µM cell strainer followed by a 40 µM cell strainer, centrifuged at 300g, resuspended in RBC cell lysis buffer (Sigma; R7757), lysis inactivated with PBS, centrifuged at 300g, and resuspended in FACS buffer. Cells were stained with anti-CD45 conjugated to APC (1:100, eBioscience, 17-0451-83), PDPN conjugated to PE-cy7 (1:200, Biolegend, 127410), and 20 ng/mL DAPI. DAPI-; CD45-; PDPN-; GFP-; tdTomato+ cells were identified as AT2 cells and sorted to purity. DAPI-, CD45-; PDPN+; GFP+; tdTomato-cells were identified as AT1 cells and sorted to purity.

### Bulk RNAseq of sorted AT2 and AT1 cells

RNA from sorted AT2 and AT1 cells was isolated using Direct-zol RNA miniprep (Zymo Research; R2050). RNA yield was quantified, libraries generated, and sequenced according to standard methods and have been previously described.^52^

### RASL-seq probe design and sample prep

The RASLseq assay was adapted from a previously published protocol.^53^ Briefly, *Hopx-*GFP murine organoids were plated in 384 well format in Matrigel® (Corning) and incubated with compound for 12-13 days. After treatment, Matrigel® was dissolved using Cell Recovery Solution (Corning) at 4C and washed in DPBS (Gibco). Cultures were then aspirated to 10uL and an equal volume of Proteinase-K RASL lysis buffer was added to the well. Samples were then processed as published.^53^ For detection of mouse specific lung epithelial markers, a panel of 127 genes was used with 3 oligo pairs designed per gene (Supp. Table 1). For balancing gene expression, the custom lung panel was run in conjunction with a 978 gene panel.^54^

### RASL-seq data processing

Count matrix was obtained by mapping the raw reads to the probe sequences (Supp. Table 1) as previously described^53^, and samples with <15,000 mapped reads were removed. Next, read counts were normalized by the total mapped reads, i.e., RPM (Reads Per Million). Out of three probes designed for each gene, the one with the most consistent measurement to a previous benchmarking RNA-seq dataset (not shown) was chosen to serve as the expression level value of that gene. Consistency of measurement was evaluated by inter-replicate similarity determined by the Pearson correlation coefficient of log2(RPM+1) of all the genes. Samples with maximum inter-replicate similarity < 0.8, or >40% undetected probes compared to the best replicate, were considered as outliers, and excluded from downstream analysis. Batch effect across plates was removed using pyCombat^55^, then quantile normalization was applied to adjust log2(RPM+1) across replicates. Differential expression analysis was performed using limma^56^ in R.^57^

### scRNA-seq data analysis

Raw reads were mapped to the mm10 mouse reference genome and a per-cell count matrix was generated using CellRanger pipeline (10X Genomics, version 5.0.1).^58^ The processed count matrix was used to filter out low quality cells with <700 detected genes or >10% mitochondrial reads. Doublets were identified using DoubletFinder and excluded from downstream analysis.^59^ UMI counts were normalized by Seurat sctransform ^60^ and top 2000 variable genes were selected for dimension reduction and 10-15 principal components were used for Leiden clustering at the downstream of Seurat v4 framework.^61^ To ensure identification of rare and transient cell types and states, we set resolution to 1.0 for initial cell clustering and merge clusters of the same type based on cluster markers identified by FindAllMarkers function in Seurat. Cells from replicates of each condition were pooled together and major cell categories including epithelial, endothelial, mesenchymal, and immune cells, were identified first using the described workflow. Cell types within each category were then identified using the same protocol.

### Data integration and batch removal

Although cells were highly consistent across replicates of the same treatment, marked transcriptomic changes were observed in cells from different conditions. To identify matched cell types and states and remove batch effect across conditions, integrated cells across conditions using canonical correlation analysis in Seurat on 2000 most variable genes.^62^ Results of data integration were inspected on UMAP with cell type labels previously identified for each condition. For comparative analysis to bleomycin treated mouse lung, mesenchymal cells from GSE132771 were integrated with our data using the same method.^11^

### Differential expression and gene set enrichment analysis

To unbiasedly identify differentially expressed genes (DEGs) induced by compound treatment, we employed a cell label agnostic approach that searches DEGs from unsupervised cell clusters. Pseudobulk-based framework was used to reduce impact of dropout and technical variability attributed to single cell sequencing.^63^ Specifically, raw UMI counts from all cells of the same type were pooled together for each biological replicate and fed into pseudoBulkDGE function of Scran package in R Bioconductor.

Differentially expressed protein coding genes with FDR<0.01 were ordered based on their log2 fold change. To identify enriched gene sets, we ran the fGSEA package v1.28.0^64^ with default setting while limiting the gene set sizes between 5 and 500 genes. Cell clusters with enriched gene sets (p<0.05) in their DEGs were selected for further analysis.

To quantify pathway activity in cells, AddModuleScore function in Seurat was used to estimate a combined expression level of all pathway genes relative to randomly chosen genes. Cell cycle score was determined in a similar way using the CellCycleScoring function and default cell cycle genes in Seurat.

### RNA velocity and trajectory analysis

Kallisto bustool pipeline (version 0.46.2)^65^ was used to align reads onto mouse genome (Ensembl assembly release 101, 3’ cDNA) using the option --fr-stranded. Spliced and unspliced reads were counted by Velocyto.^19^ Data were then normalized by Scanpy^66^ and RNA velocity was calculated by scVelo package.^67^ Moments were calculated using the scvelo.pp.moments() function with parameters pcs = 15 and neighbours = 10. Velocity estimation was performed using the scvelo.tl.velocity() function with default stochastic mode. To estimate trajectory of AT2 cells differentiating into *Fn1*+ and *Fn1*-*Krt8*+ ADI cells, diffusion map was calculated by Destiny package^68^ with 2000 most variable genes and 15 pcs.

### Cross species comparison

Similarity of human and mouse lung epithelial cell types was estimated using highly expressed genes. Specifically, single cell data from IPF patients (GSE135893) was reprocessed according to the published protocol.^7^ Next, average gene expression for each cell type was calculated by AverageExpression function in Seurat and low expressing genes with max log expression <1 across all cell types were excluded. Finally, ortholog genes from two species with same symbols were selected and the Spearman correlation coefficient was calculated for each pair of cell types.

### Experimental design, statistical analysis and plotting

EC50 was calculated on log transformed compound concentrations using a three-parameter dose-response curve (Domatics, Graphpad Prism). Zero concentration controls were set as suitably low dose to not adversely effect results. All graphical figures were created with BioRender.com. All figures were assembled in Illustrator 2023 (Adobe). Plots were generated in Prism 9 (Domatics, Graphpad Prism).

## ACKNOWLEDGEMENTS

We thank D. Quackenbush and F. Lo for HTS imaging and analysis; M. Lindstrom and vivarium staff for animal caretaking; C. Trussell for flow sorting; and F. Luna for RNA sequencing.

## DATA AVAILABILITY

scRNA-seq data generated in this study will be uploaded to the Gene Expression Omnibus (GEO).

## AUTHOR CONTRIBUTIONS

Conceptualization: D.P.P, B.F., R.S.D.; Investigation: S.W., C.J.N.M, S.Y., B.N., G.C.F., G.K., J.C.S., J.H., V.C., S.W.B.; Methodology: F.J.K., B.T., J.R.W., D.P.P., B.F., R.S.D.; Formal analysis: S.W., C.J.N.M., S.Y., B.N, G.C.F., G.K., J.C.S., J.H., V.C., S.W.B., F.J.K., B.T., J.R.W., D.P.P., B.F., R.S.D.; Visualization: A.S.H., B.F., G.K.; Writing - original draft: A.S.H., B.F., R.S.D.; Writing - review & editing: A.S.H., S.W., C.J.N.M., S.Y., B.N., G.C.F., G.K., J.C.S., J.H., V.C., S.W.B., F.J.K., J.R.W., J.E., D.P.P., B.F., R.S.D.; Supervision: F.J.K., B.T., J.R.W., J.E., D.P.P., B.F., R.S.D.

---

**Supplementary Figure 1:**
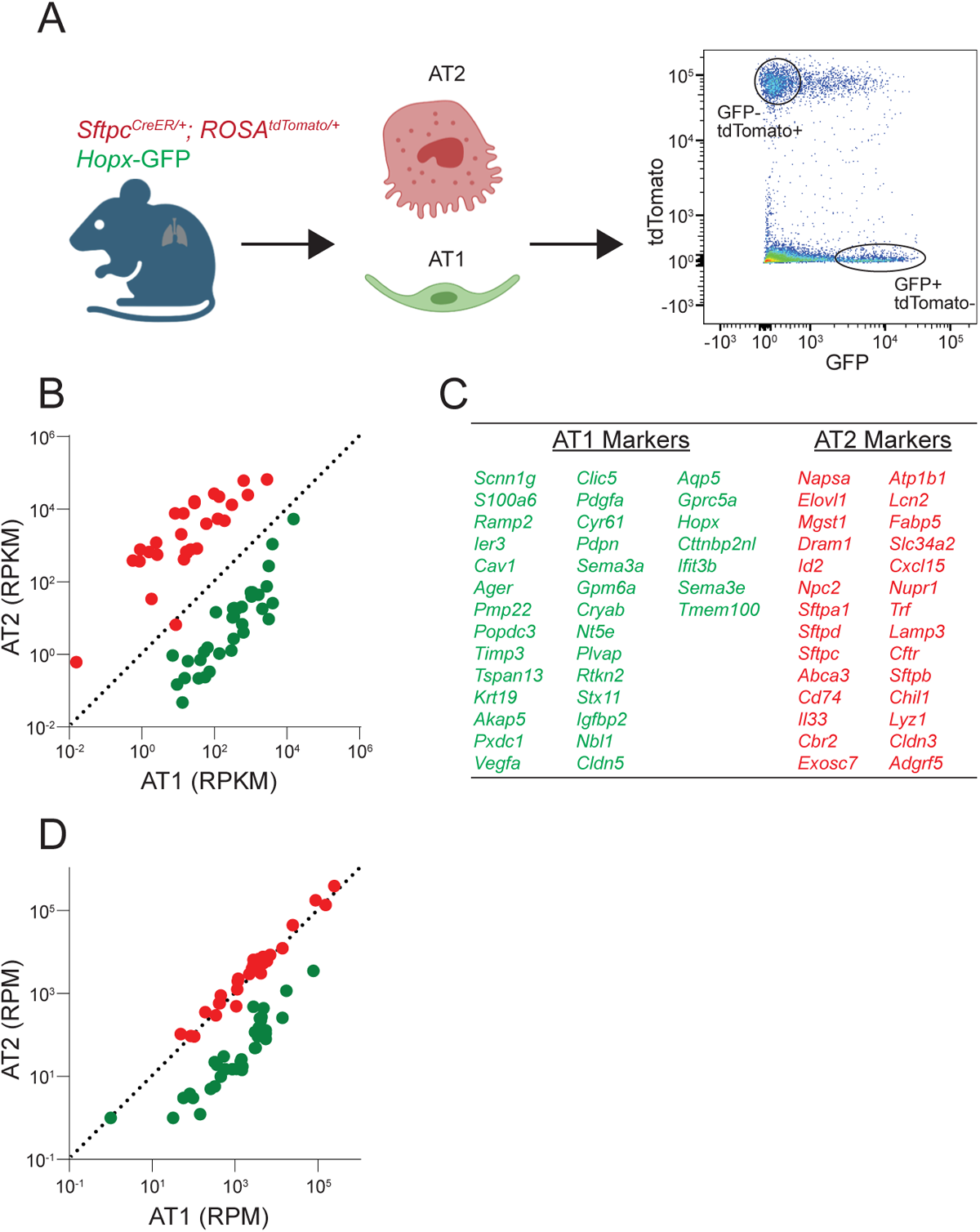
Development of a custom RASL panel enriched for AT2 and AT1 cell lineage markers. **A**: Schematic of assay. Whole distal lung cells from adult, tamoxifen induced SftpcCreER;ROSA26*^tdTomato^; Hopx-GFP* mice were sorted into tdTomato+ and GFP+ populations for bulk RNA sequencing. **B**: Expression of selected genes that were enriched in GFP+ population (AT1) vs tdTomato+ population (AT2). Full expression profiles on samples listed in Table 1. RFPM= reads per kilobase per million. Dotted line is y=x+0. **C**: List of markers identified as specific for AT1 or AT2 cells. **D**: Expression of RASL probes targeting selected genes confirms enrichment in GFP+ population (AT1) vs tdTomato+ population (AT2). RPM=reads per million. Dotted line is y=x+0.

**Supplementary Figure 2:**
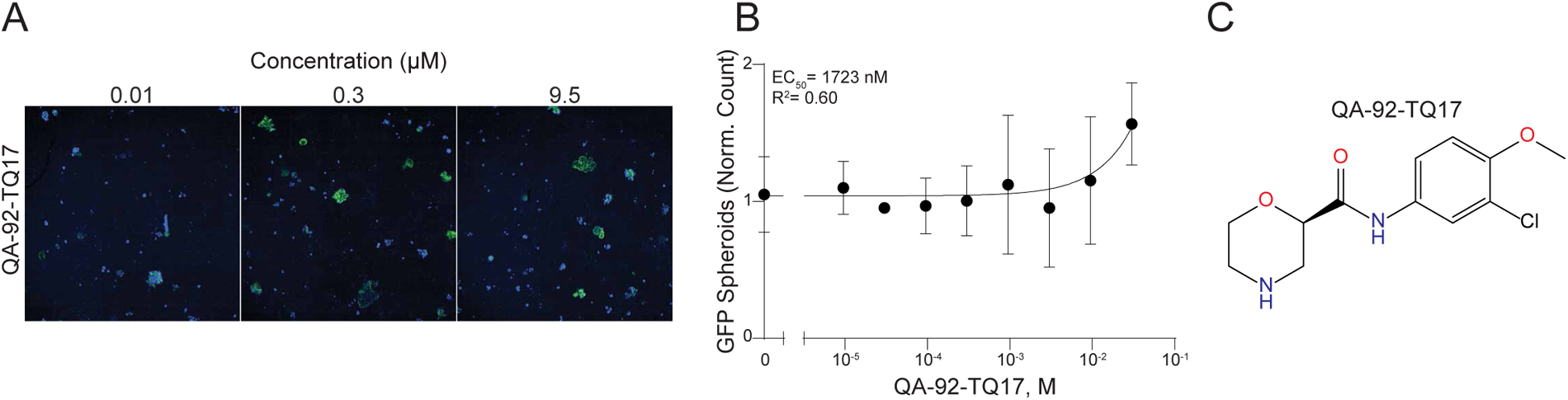
DB-11-BE87 analogue, QA-92-TQ17, does not rescue Tgfβ impaired alveolar type 1 spheroid formation. **A**: Representative images of live alveolospheres treated at time of plating with increasing doses of QA-92-TQ17. **B**: Quantification of concentration-response curve for QA-92-TQ17 on GFP adjusted count spheroids. EC_50_= 1723 nM, R^2^=0.60. n=2 for all concentration except untreated well, n=48 for untreated well. mean ± SD **C**: Chemical structure of QA-92-TQ17.

**Supplementary Figure 3:**
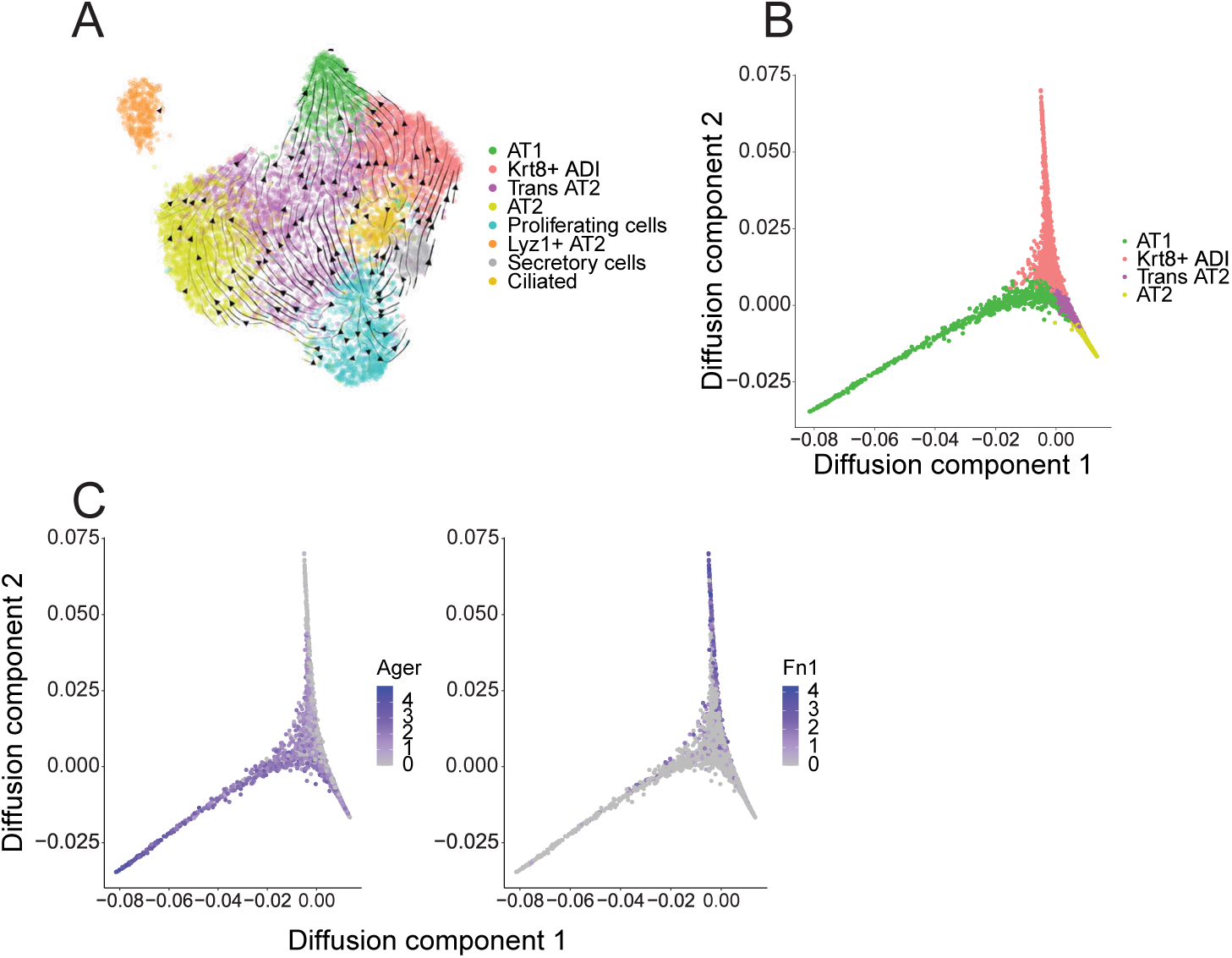
Krt8+ ADI and AT1s are separate terminal cell fate branches. **A**: RNA velocity vectors projected over UMAP plot of epithelial subclusters integrated from all samples. **B**: Diffusion map of AT1, AT2, *Krt8*+ ADI, and Transitional AT2 epithelial cell types. **C**: Expression of individual genes on diffusion map from B: *Ager* (left) and *Fn1* (right).

**Supplementary Figure 4:**
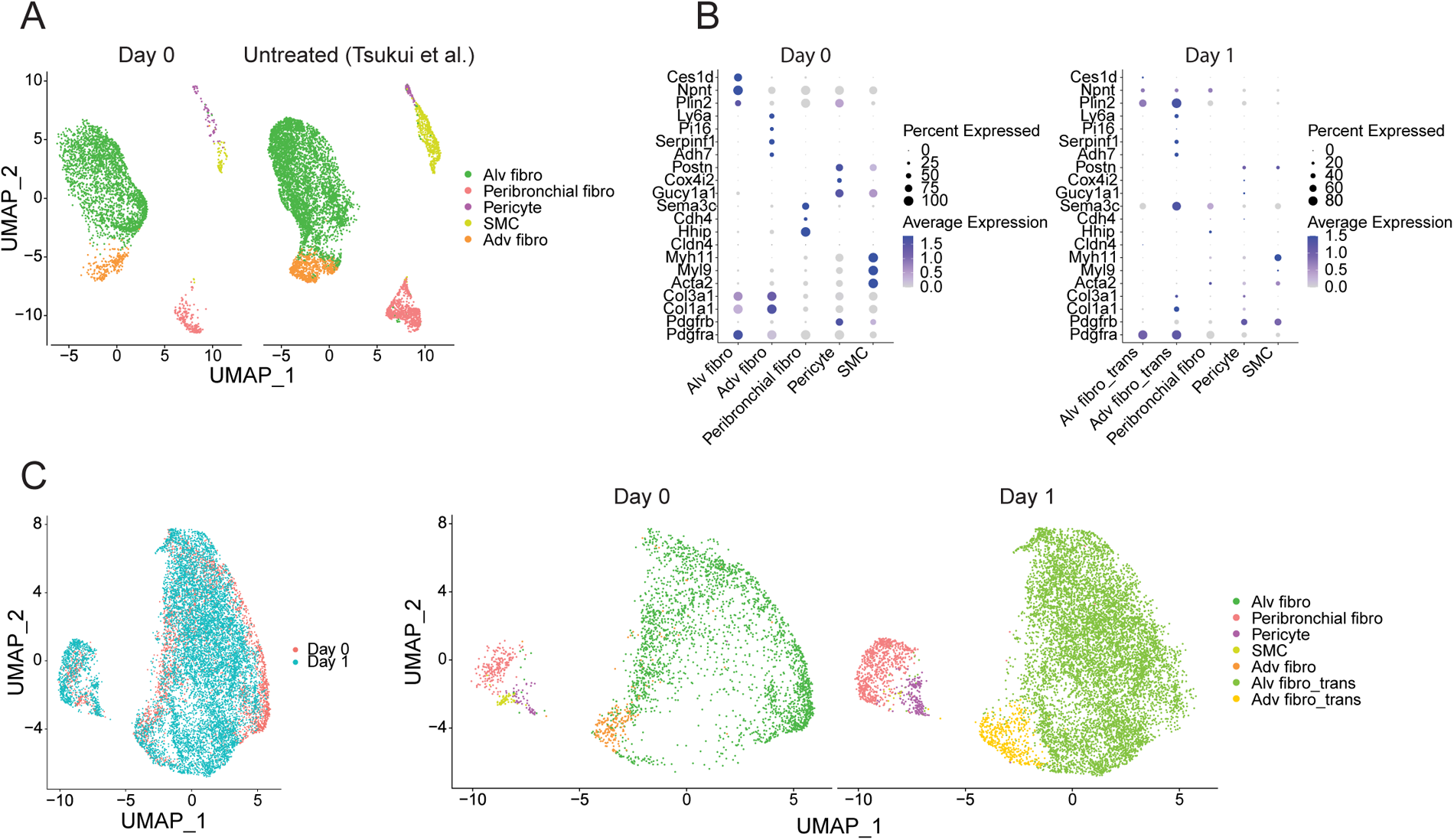
Mesenchymal cells in alveolosphere assay shift cellular identity at early timepoint. **A**: UMAP plot of integrated mesenchymal cells from two studies. Left, cell mixture added to culture on day 0. Right, untreated mouse lung from Tsukui, T. *et al.,* colored by cell type labels from publication. **B**: Dot plots of marker expression in mesenchymal subtypes, split by collection timepoint (day 0, left, and day 1, right). **C**: UMAP plot of integrated mesenchymal cells from day 0 (red) and day 1 (teal). Cell types are indicated with colors in UMAPs split by time points (day 0, middle, and day 1, right).

**Supplementary Figure 5:**
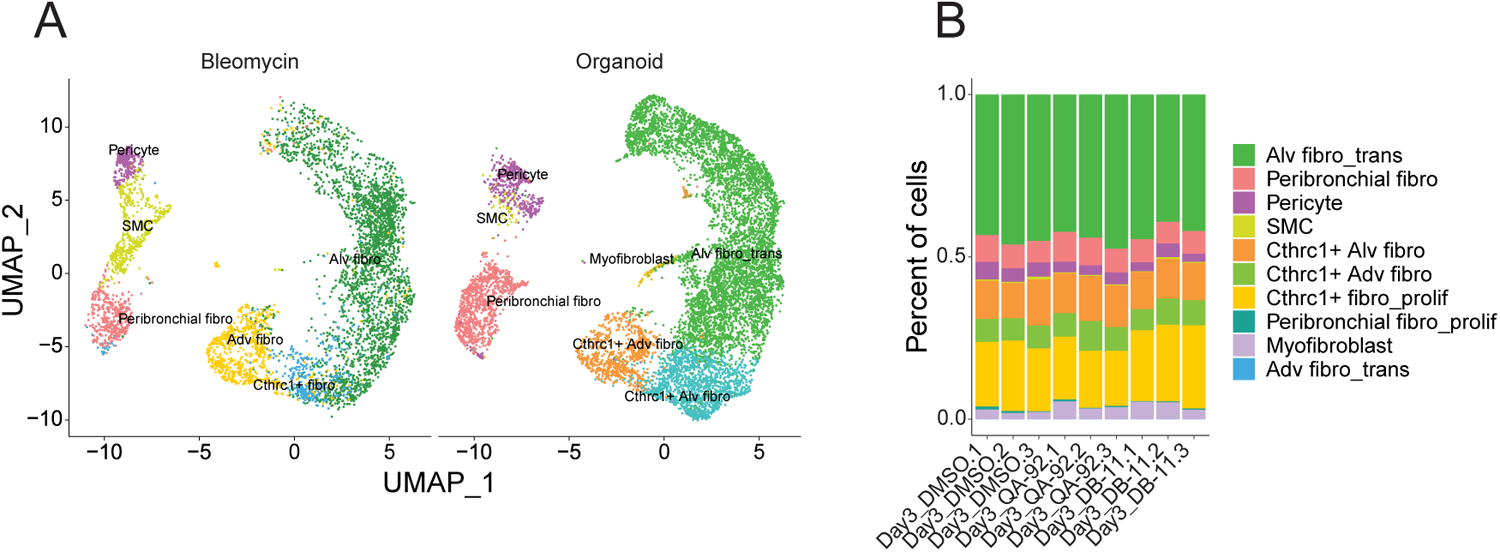
Mesenchymal subtype proportion did not change at the day 3 timepoint. **A**: UMAP plot of integrated mesenchymal cells from two studies. Right, alveolar organoids on day 3. Left, bleomycin treated mouse lungs (Tsukui, T. *et al.*), colored by published cell type labels. **B**: Proportion of mesenchymal subtypes from day 3 samples, separated by compound treatment.

**Supplementary Figure 6:**
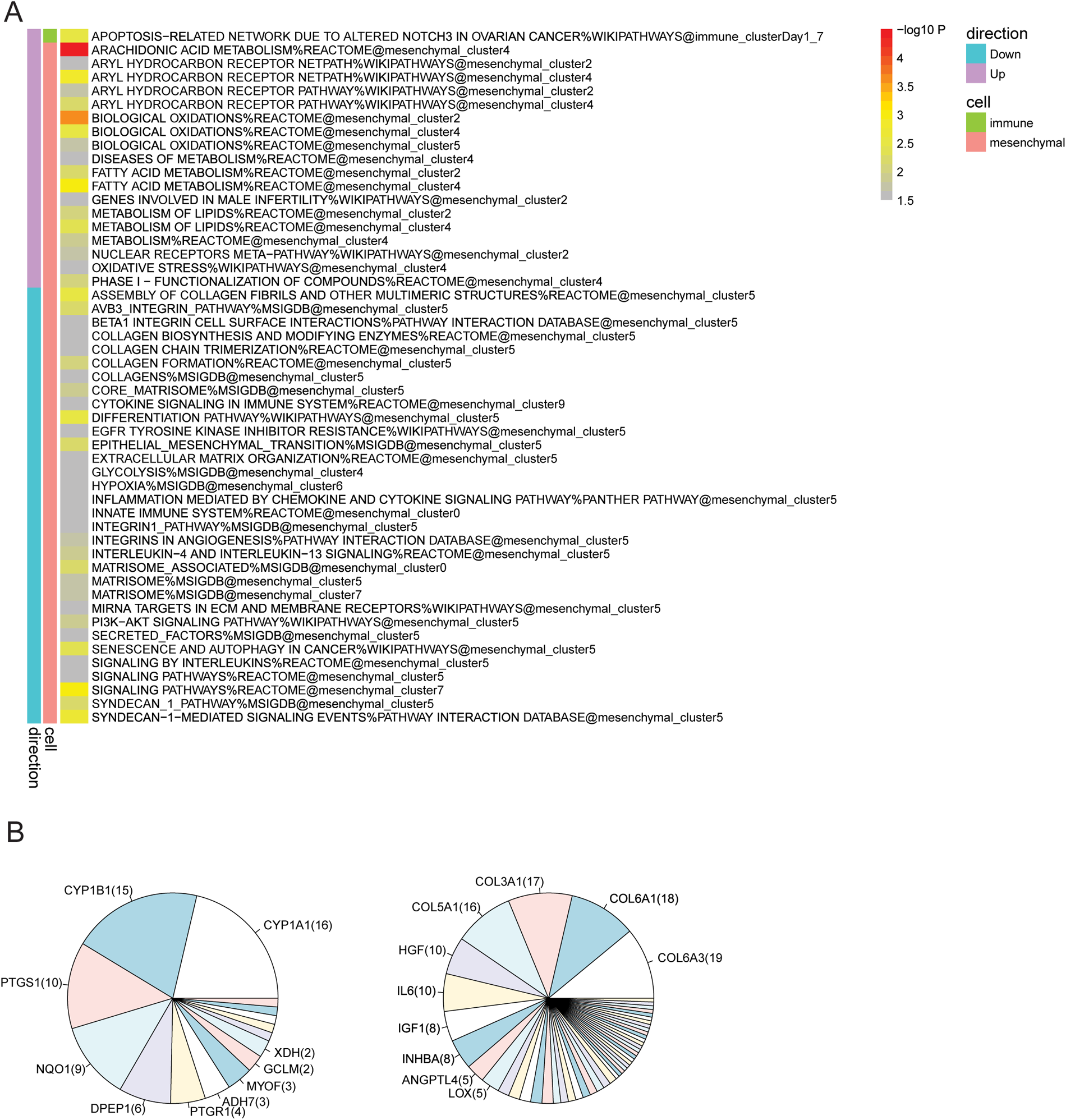
DB-11-BE87 activates AHR pathway early. **A**: Heatmap of pathways and biological processes enriched for differentially expressed genes (DB-11-BE87 vs. DMSO) across all cell clusters in samples from day 1. Left bar indicates direction of DEGs (down, teal; up, purple). Center bar indicates cell category (immune, green; mesenchymal; pink). Right bar indicates degree of enrichment of DEGs. Cell clusters without significant enrichment were excluded. **B**: Pie chart of most frequently regulated genes across immune and mesenchymal subclusters (left, up-regulated; right, down-regulated).

**Supplementary Figure 7:**
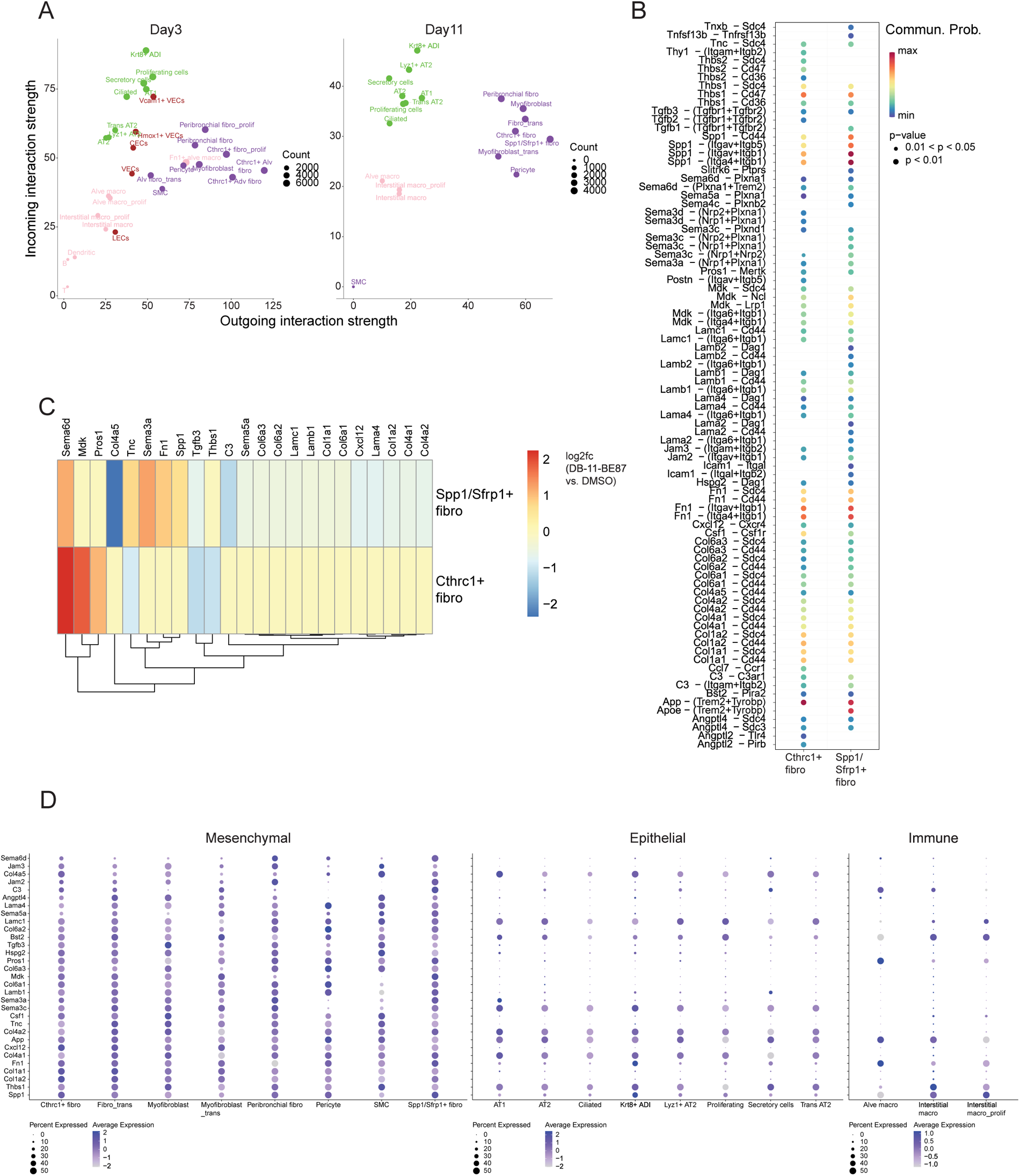
Cell-cell communications between macrophages and fibroblasts. **A**: Scatter plots of the comparison between overall strength of out-going and incoming interactions for each cell type, from DB-11-BE87-treated samples on day 3 and day 11. Size of dot indicates total cell count. **B**: Predicted ligand-receptor pairs mediating communications going from two fibroblast subtypes into interstitial macrophages, from DB-11-BE87-treated samples on day 11. Color indicates probability of each interaction and size of dot indicates statistical significance. **C**: Heatmap of DB-11-BE87-induced expression changes on 23 out of 31 ligands mediating common interactions in B. Ligands unaffected by DB-11-BE87 in both cell subtypes were excluded. Colors indicate log2 fold changes on day 11. **D**: Dot plot for expression levels of 31 ligands mediating common interactions in B, across all cell types in the DB-11-BE87-treated samples on day 11.

